# Encoding of odor information and reward anticipation in anterior cortical amygdaloid nucleus

**DOI:** 10.1101/2020.11.19.390740

**Authors:** Kazuki Shiotani, Yuta Tanisumi, Junya Hirokawa, Yoshio Sakurai, Hiroyuki Manabe

## Abstract

Olfactory information directly reaches the amygdala through the olfactory cortex, without the involvement of thalamic areas, unlike other sensory systems. The anterior cortical amygdaloid nucleus (ACo) is one of the olfactory cortices that receives olfactory sensory input, and is part of the olfactory cortical amygdala, which relays olfactory information to the amygdala. To examine its electrophysiological features, we recorded individual ACo neurons during the odor-guided go/no-go task to obtain a water reward. Many ACo neurons exhibited odor-evoked go cue-preferred during the late phase of odor-sampling supporting the population dynamics that differentiate go/no-go responses before executing the odor-evoked behaviors. We observed two types of neurons with different anticipation signals: one neuron type exhibited gradual increases of activity toward reward delivery, while another type exhibited a phasic go cue-preferred activity during odor sampling as well as another phasic anticipatory activity for rewards. These results suggest that the ACo may be involved in reward-related behavioral learning by associating the olfactory information with reward anticipation.

## Introduction

Olfaction is closely related to emotion in attributing positive (attractive) or negative (aversive) valence to the environment more than any other sensory modality ^12^. The close bidirectional connection and the particular organization of the olfactory cortex to the amygdala distinguishes the olfactory system from other sensory systems ^34^. Afferent sensory inputs from the main olfactory bulb (OB) directly target the amygdala through the olfactory cortex, while afferent inputs from most of the other sensory systems enter the amygdala via the thalamus and neocortical regions ^5^. OB mitral cells project their axons through the lateral olfactory tract to the olfactory cortex ^6^. Olfactory cortical amygdala, which is a part of the olfactory cortex, relays olfactory information to the amygdala. The anterior cortical amygdaloid nucleus (ACo) is a part of the olfactory cortical amygdala, and has a bidirectional connection with the amygdala.

A study reported that the ACo receives dense projections from the main olfactory bulb (OB), moderate projections from the piriform cortex, lateral entorhinal cortex, basomedial amygdaloid nucleus (BMA), and medial amygdaloid nucleus (Me), and scarce projections from the ventral tegmental area (VTA) and the ventral tenia tecta (vTT) ^3^. Moreover, the ACo projects densely to BMA ^3^. These anatomical studies indicate that the ACo is closely related to the amygdala, and it is possible that the ACo is involved in odor-evoked motivational behaviors.

A behavioral study revealed that ACo participates in olfactory fear conditioning in rats as electrical stimulation of the olfactory bulb induces evoked field potential signals (EFPs), that are persistently potentiated specifically in the ACo after training ^7^. Moreover, electrical stimulation of the ventral tegmental area (VTA) showed that the ACo, besides other mesolimbic structures, displays increased Fos expression in rats ^8^. A whole-cell patch clamp study showed that with the activation of sodium conductance, pyramidal neurons of the ACo displayed rhythmic fluctuations of intrinsically generated voltage-dependent membrane potential in the theta-low beta range, suggesting that the ACo was related to synaptic plasticity and learning ^9^. ACo has been poorly investigated, but comprehensive evidence suggests that it may play a prominent role in reward-related behavioral learning by olfactory stimulation. However, little is known about the electrophysiological features of the ACo neurons for reward-related behavioral tasks.

Here, we recorded the neural activity of ACo neurons during odor-guided reward-directed behaviors. Many ACo neurons responded to the go-cue odor stimulus at the late phase of the odor-sampling epoch (from the odor valve off to the odor port exit). The ACo neuron population showed profound and persistent transformations in the dynamics of cue encoding over 400 ms after odor onset. Furthermore, we found that the ACo neuron groups each coded a different type of anticipation signal: one neuron group type exhibited gradual increases in the signals to the reward, while the other type showed phasic anticipation signals with the go-cue preference responses during odor sampling. Our results suggest that the ACo neurons may play an important role in odor-guided reward-directed learning.

## Results

### Go-Cue-Odor preferred responses of ACo neurons during the late phase of odor-sampling epoch

We recorded 158 well-isolated neurons in the ACo of four mice performing an odor-guided go/no-go task (Fig. 1a, recording positions are shown in Fig. 1b). Briefly, the go trial requires the mice to first sample a go-cue odor presented at an odor port and then to move to a reward port to receive water reward. Conversely, the no-go trial requires the mice to first sample a no-go-cue odor presented at the odor port and then to stay near it to wait for the next trial. It is important to note that the mice were required to keep their nose inserted into the odor port, at least during odor presentation (500 ms). After the mice were well trained, their behavioral accuracy remained above 80% throughout the session. For all mice, the median duration of the odor-sampling epoch (the time from odor valve opening until the mouse withdrew its snout from the odor port) was 1053 ms (interquartile range: 902–1212 ms) in the go trials, and 764 ms (interquartile range: 657–968 ms) in the no-go trials (31 sessions from four mice).

**Fig. 1.**
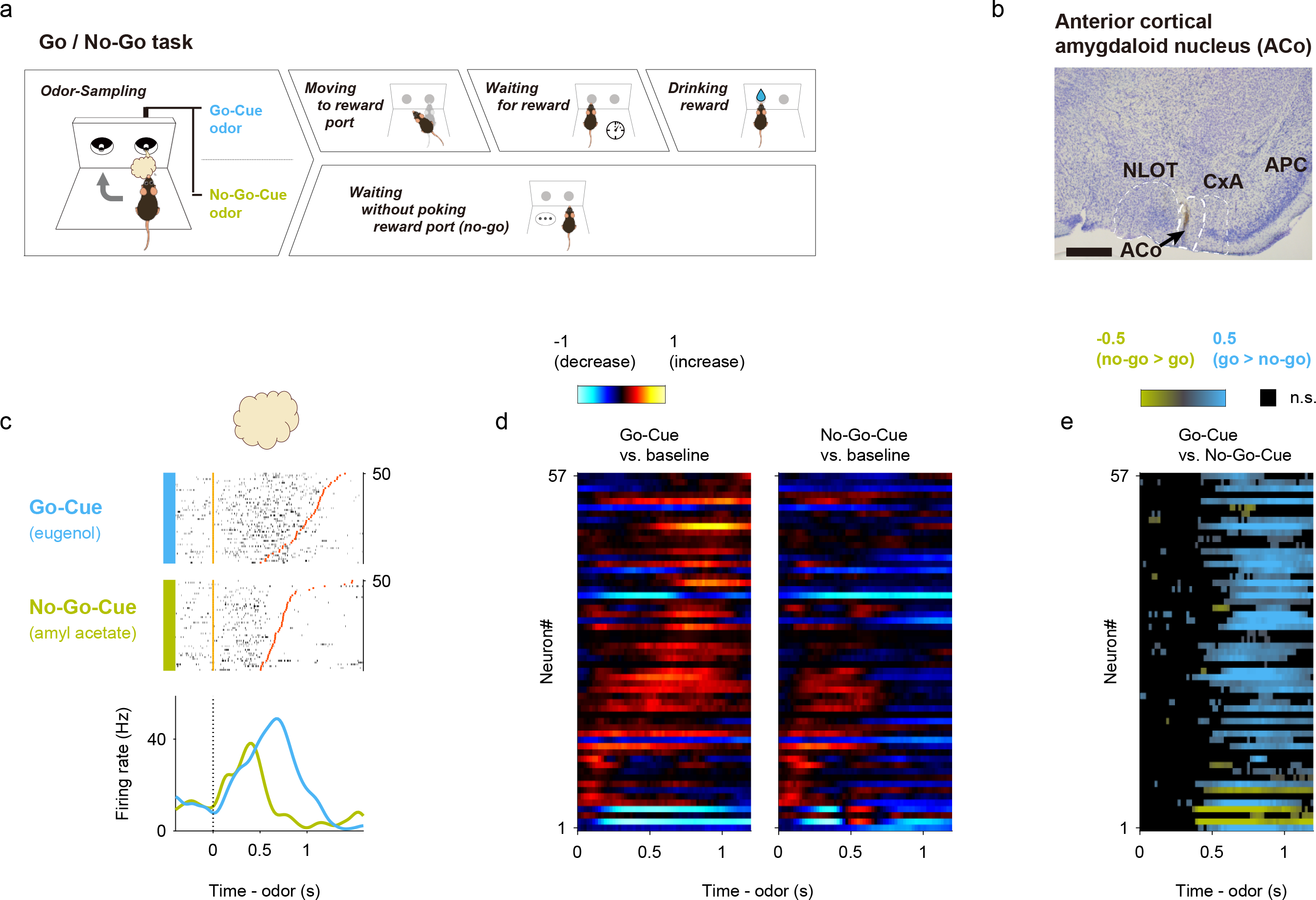
Cue-Odor-preferred responses of ACo neurons during the late phase of odor-sampling in the odor-guided go/no-go task. (**a**) Time course of the odor-guided go/no-go task. Behavioral epoch temporal progression from left to right. (**b**) Nissl-stained frontal section (an arrow indicates recording track) and recording tracks (vertical thick lines) of the ACo. NLOT, nucleus of lateral olfactory tract. CxA, cortex-amygdala transition zone. APC, anterior piriform cortex. Scale bar: 500 μm. (**c**) Example firing patterns of ACo neurons during odor-sampling epoch (the time from odor valve opening to odor port exit) in the odor-guided go/no-go task. Each row contains spikes (black ticks) for one trial, aligned to the time of odor valve opening (corresponding to odor port entry, orange ticks). Red ticks represent times of odor port exit. Correct trials are grouped by odor, and within each group, are sorted by the duration of the odor-sampling epoch (50 selected trials from the end of the session are shown per category). Histograms are averaged across odors, and calculated using a 20 ms bin width and smoothed by convolving spike trains with a 60 msec-wide Gaussian filter (blue, go-cue odor; green, no-go-cue odor). Vertical dashed lines indicate the time of odor valve opening. (**d**) Normalized firing rates (auROC values) for go-cue selective neurons (n = 57). auROC values (aligned by odor valve opening) were calculated by go-cue odor presentation versus baseline (left) and no-go-cue odor presentation versus baseline (right) in the sliding bins (width, 100 ms; step, 20 ms). Red, increase from baseline; blue, decrease from baseline. Each row corresponds to one neuron, with neurons in the left and right graphs in the same order. Neurons are sorted by the peak time for auROC values calculated by go-cue odor presentation versus baseline. (**e**) Cue preference curves (auROC values, go-cue versus no-go-cue odor presentation, aligned by odor valve opening, odor port exit) for go-cue selective neurons. Each row corresponds to one neuron, with neurons in the left and right graphs in the same order of (d). Color scale indicates significant preferences (p < 0.01, permutation test; positive values correspond to the go-cue preferred responses). The black boxes indicate bins with non-significant preferences (p > 0.01, permutation test).

Since the ACo receives direct inputs from the mitral cells of the olfactory bulb, we first focused on whether ACo neurons exhibited cue-odor selective activity during odor-sampling epochs (from odor poke-in to odor poke-out). We found that a subset of ACo neurons increased their firing rates during the odor presentation phase (0–500 ms after the odor valve opening) during both go and no-go trials, and then showed a go-cue-odor preferred response 500 ms after the odor onset (an example shown in Fig. 1c). To quantify the dynamics of the cue-encoding, we calculated the firing rate changes from baseline (200 to 0 ms before the end of the inter-trial interval) in the sliding bins during the odor-sampling epoch for each neuron. For each correct trial, we calculated the area under the receiver operating characteristic curve (auROC) value at each time bin (width: 100 ms, step: 20 ms), and defined the go-cue selective neurons (n = 57 neurons, 36.1 % of the recorded neurons) as those neurons that significantly increased their firing rates from the baseline (p < 0.01, permutation test) for five consecutive bins (100 ms) during the odor-sampling epoch in the go correct trials (Fig. 1d). Across the go-cue odor-selective population, calculation of go-cue versus no-go-cue preferences during odor-sampling epochs clearly showed a go-cue preference manner from 500 ms after the odor onset to the odor poke-out (late phase of the odor sampling epoch) (Fig. 1e, p < 0.01, permutation test). These results suggest that the ACo received not only a particular odorant profile directly from the olfactory bulb but rather the complex odor information, including behavioral contexts from other olfactory cortical areas and top-down inputs from higher areas.

### Late phase of go-cue odor preferred responses were evoked by the odor onsets and were stable across trials

The go-cue odor-selective population showed cue-odor-preferred responses during the late phase of the odor sampling epoch (Fig. 1c). It is possible that the late phase of odor-preferred responses was tuned to the odor port exit behaviors or contained the premotor signals that were observed in many brain regions ^10,11,12^. To take these signals into account, and to help isolate signals related to odor presentation and action, we developed an encoding model (generalized linear model, GLM). This model incorporated task-related variables during the odor-sampling epoch as predictors of each neuron’s activity (Fig. 2a and Supplementary Figs. 1a-c) ^13^.

**Fig. 2.**
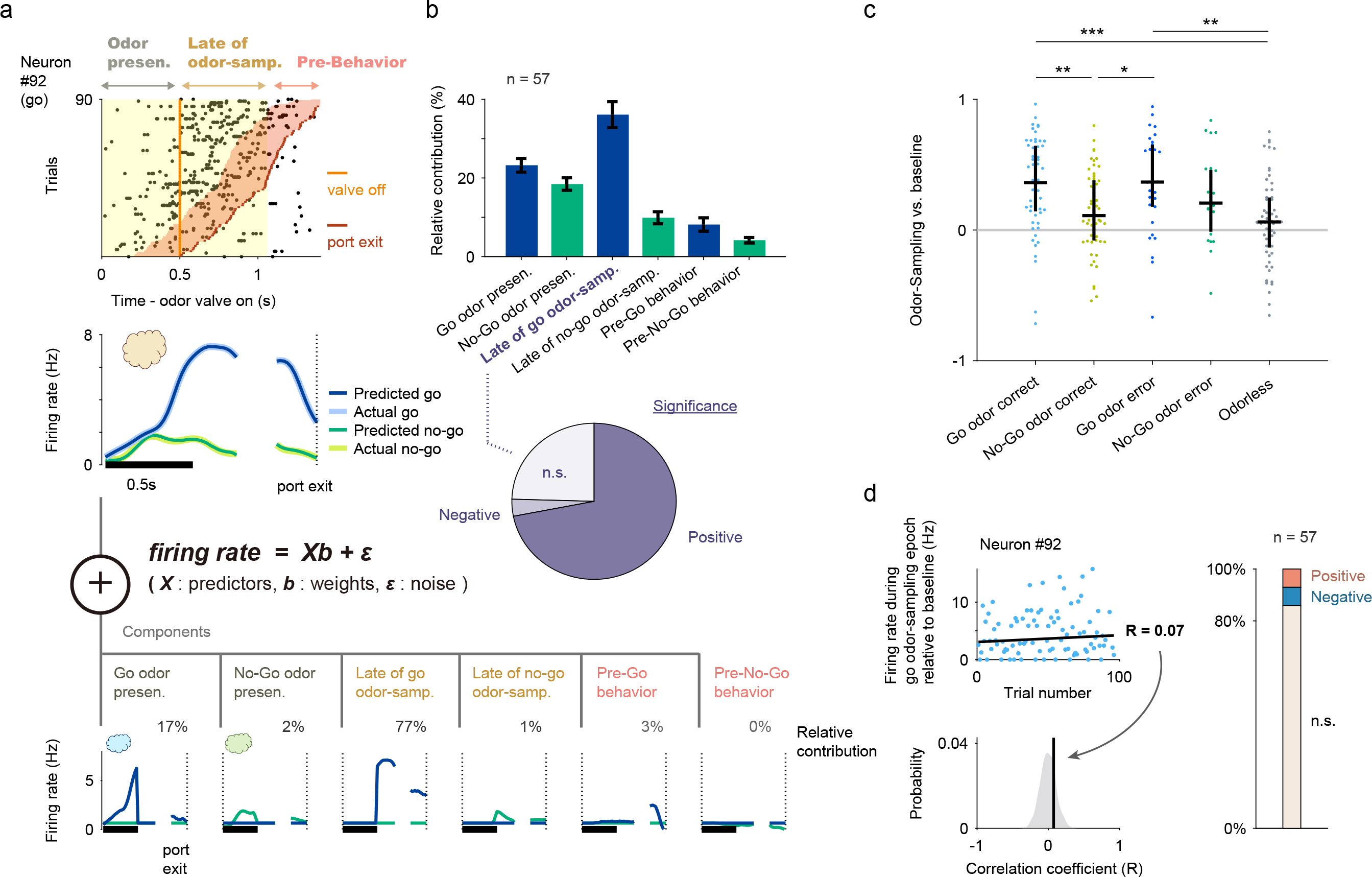
Late go-cue odor-preferred responses were evoked by the odor onsets and were stable. (**a**) Schematic of the encoding model used to quantify the relationship between behavioral variables and the activity of each neuron (see Materials and methods). Behavioral predictors for the odor stimulus-presentation epoch are supported over the window 0 to 500 ms relative to the onset of odor valve, as well as late phase of odor-sampling epoch that are supported over the window 0 to 553/264 ms relative to the offset of odor valve in either go/no-go trials (median of the valve offset to odor port exit), and pre odor port exit epoch that are supported over the window -300 to 0 ms relative to the odor port exit. Inset, predicted and actual averaged firing rate relative to the odor onset and odor port exit for one neuron. (**b**) Top: relative contribution of each behavioral variable to the explained variance of the neural activity, averaged across the go-cue-selective neurons. All error bars represent the standard error of the mean. Bottom: relative contribution significance of the late phase of go-cue odor-sampling variable; see Supplementary Fig. 1f for the other variables. (**c**) Go-cue odor-preferred responses during correct trials, error trials, and catch (odorless) trials. The auROC values were calculated during the odor-sampling epochs and only neurons with a minimum number of three trials for each analyzed condition were included in this analysis. Black horizontal lines and black vertical lines indicate medians and interquartile ranges. The statistical significance among five groups (*p < 0.05, **p < 0.01, ***p < 0.001) was assessed by one-way analysis of variance (ANOVA) with Tukey’s post hoc test. (**d**) The development of go-cue responses in go-cue-selective neurons during learning. For each go-cue-selective neuron, we calculated the correlation between the firing rate during the go-cue odor-sampling epoch relative to the baseline (a mean firing rate during inter trial interval was subtracted for each neuron) and the order of go trial from the start of the session. The correlation coefficient was compared with control values calculated by the 1000 trial-shuffled data (gray shaded area) and then the statistical significance was determined (< 0.5^th^ percentiles of the control values, negative correlation; > 99.5^th^ percentiles of the control values, positive correlation). Across go-cue-selective neurons, the majority of the go-cue responses were not correlated with trial progression (86.0%, not significant; 7.0%, negative; 7.0%, positive).

Using this encoding model, we quantified the relative contribution of each behavioral variable during the odor-sampling epoch to the response of each neuron by determining how much the explained variance declined when that variable was removed from the model (see Materials and methods; a relative contribution for an example neuron is shown in Fig. 2a and Supplementary Fig. 1e). Averaged across the go-cue odor selective population, the highest relative contribution during odor-sampling epochs was attributed to late go-cue odor sampling (36.1 ± 3.3% of the total variance explained during the odor-sampling epoch), followed in descending order by the go-cue odor presentation (23.2 ± 1.7%), the no-go-cue odor presentation (18.5 ± 1.6%), the late no-go-cue odor sampling (9.9 ± 1.5%), pre-go-behavior (8.2 ± 1.7%), and pre-no-go behavior (4.2 ± 0.7%) (bars in Fig. 2b). The relative contributions of the late go-cue odor sampling were significantly positive across 71.9% of the go-cue odor-selective neurons (a pie chart in Fig. 2b). Furthermore, across the population, the go-cue responses during odor-sampling epochs in both correct and error trials were higher than those in the no-go-cue correct and odorless trials (Fig. 2c), suggesting that the go-cue excitation responses mainly reflected signals of encoding cue-odor information. Notably, the intensities of the majority of the go-cue responses remained stable across trials (Fig. 2d). Taken together, the go-cue-preferred responses during the late phase of the odor-sampling epoch were considered to reflect the go-cue odor information.

### Response dynamics of the ACo neuron population during the late phase of odor-sampling epoch

We demonstrated that ACo neurons showed odor-evoked cue-preferred responses during the late phase of the odor-sampling epoch (Figs. 1-2). Were the distinct cue responses reflected in the ACo neuron population dynamics, and how much could the population activity account for animals’ behavioral accuracy? First, to gain insight into the dynamics of the population response, we visualized average population activity using principal component analysis, a dimensionality reduction method. Fig. 3a shows trajectories of the mean response of the ACo neuron population to go-cue and no-go-cue odors, represented as projections onto the first three principal components (PC) during the odor-sampling epochs. Throughout the approximately 400 ms interval from the odor onset, trajectories remained converged, showing little difference across conditions. Over the late phase of odor-sampling epochs, trajectories in the odor-sampling epoch subspace began to spread out and were clearly separated at the population level. To quantify these observations, we measured the instantaneous separation between the population cue responses (Fig. 3b). The separation started to increase from 400 ms after odor onset, reaching a maximum at ∼800 ms, and remained above baseline levels until odor port exit. Thus, the ACo neuron population showed profound and persistent transformations in the dynamics of cue-encoding, 400 ms after odor onset.

**Fig. 3.**
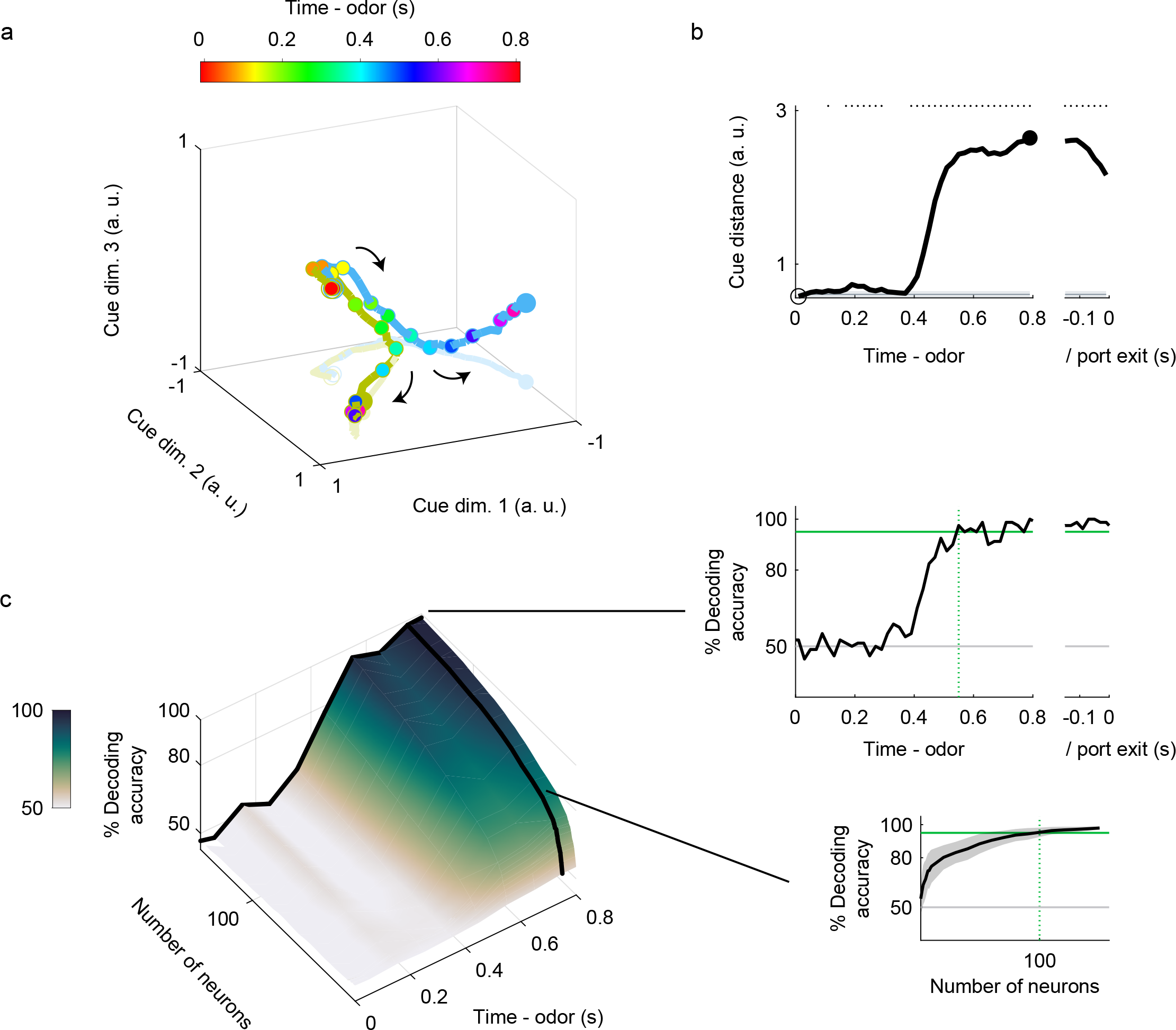
Dynamics of ACo neuron population response during the late phase of odor-sampling. (a) Visualization of ACo neuron population responses during odor-sampling epoch using principal component analysis (158 neurons). The responses to cue odors are projected onto the first three principal components corresponding to odor-sampling epoch subspaces. Blue line, go-cue odor; green line, no-go-cue odor. Temporal progression from unfilled blue/green spheres to filled spheres. (**b**) Distance between ACo neuron population responses. Gray line and shaded area show mean ± 2SD baseline values during pre-odor-sampling epoch. Top dots indicate time bins showing values more than mean + 2SD baseline values. (**c**) The time course of odor decoding accuracy. A vector consisting of instantaneous spike counts for 1–158 neurons in a sliding window (width, 100 ms; step, 20 ms) was used as input for the classifier. Training of the classifier and testing were done at every time point. Green horizontal lines indicate the level of animal behavioral performance. Gray horizontal lines indicate chance level (50%). Green vertical dashed lines indicate the first points at which the decoding accuracy reached the level of animal behavioral performance. Shaded areas represent ± SD.

Second, to examine whether the population activity accounted for the animals’ behavioral accuracy, we performed a decoding analysis to determine whether the firing rates of the ACo neuron populations could be used to classify each individual trial as go or no-go. We used SVMs with linear kernels as a decoder. Based on ACo neurons, analyses of the decoding time course, using a sliding time window, revealed that decoding accuracy was first maintained at chance levels 400 ms after the odor onset, and then increased above the behavioral accuracy level of the animals around 500–600 ms after odor onset (the top-right graph in Fig. 3c). In the 700–800 ms period, about 100 neurons provided sufficient information to account for the behavioral accuracy (the bottom right graph in Fig. 3c). Thus, a hundred ACo neurons accounted for animals’ behavioral accuracy in the late phase of odor sampling.

### Two types of reward anticipation responses of ACo neurons

We then focused on the ACo activity during odor-evoked behaviors after an odor-sampling epoch. A subset of ACo neurons gradually increased their firing rates from the time of water port entry till the reward was received, and another subset of neurons increased their firing rates while waiting for reward (examples shown in Fig. 4a). We quantified the data by calculating firing rate changes from baseline (spike data were aligned to the water port entry), and three measures from the values: “time of center of mass”, “onset time”, and “duration” (from water port entry to 1000 ms after opening the water valve, Fig. 4b, see Materials and methods). The drinking epoch selective neurons (n = 30, 19.0 % of the recorded neurons) were defined as neurons that had the time of center of mass during the drinking epoch, and the waiting epoch selective neurons (n = 14, 8.9 % of the recorded neurons) were defined as neurons that had the time of center of mass during the waiting epoch. Across the population, the drinking-epoch-selective neurons gradually increased their firing rates -190 ms before the water valve opened for 432 ms, and the waiting-epoch-selective neurons increased their firing rates 10 ms after water port entry for 108 ms (Figs. 4c-d, p < 0.01, permutation test). Thus, ACo neurons exhibited two distinct types of reward anticipation responses.

**Fig. 4.**
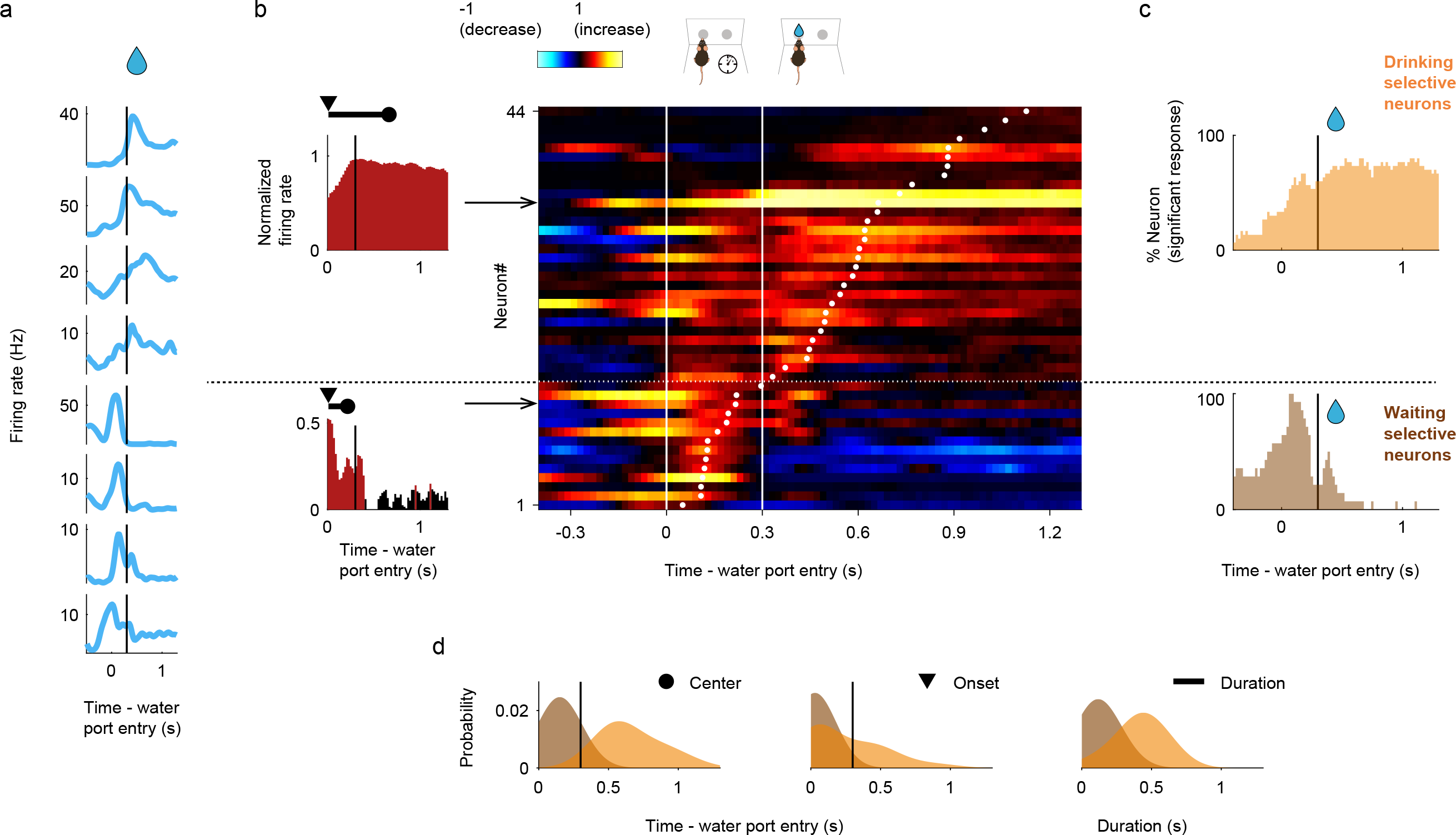
Two types of reward-related responses of ACo neurons. (**a**) Example firing patterns of reward-related responses. Spike histograms are calculated using a 20 ms bin width and smoothed by convolving spike trains with a 60 ms wide Gaussian filter. A vertical black line indicates the water valve opening. (**b**) Evaluation of the reward-related responses. Normalized firing rates (auROC values) were calculated by go-behavior versus baseline in the sliding bins (width, 100 ms; step, 20 ms). Left: red bars show significant excitation (p < 0.01, permutation test). Based on the significant time points, onset time (black triangle), time of center of mass (black circle) and duration (black horizontal line) were calculated. Vertical black lines indicate the water valve opening. Right: each row corresponds to one neuron and neurons are sorted by times of center of mass (white dots) of auROC values. Based on the times of center of mass, drinking-selective neurons and waiting-selective neurons were defined (a horizontal dashed line). Color scale as in Fig. 1d. Vertical white lines indicate the water port entry and the water valve opening. (**c**) The proportions of neurons that exhibited a significant response, calculated from auROC values (p < 0.01, permutation test) for each neuron group (orange, drinking selective neurons; brown, waiting selective neurons). Vertical black lines indicate the water valve opening. (**d**) Distributions of the times of center of mass, onset times and durations (orange, drinking-selective neurons; brown, waiting-selective neurons).

### Association of go-cue excitations with excitatory responses for the reward anticipation behavior

We observed that the waiting-epoch-selective neurons showed go-cue-preferred activity during the odor-sampling epoch; however, the drinking-epoch-selective neurons did not (examples shown in Fig. 5a). To examine the relationship between the reward anticipation responses and cue encoding, we quantified the response profiles of each neuron group during odor-evoked behaviors by calculating the firing rate changes from baseline (Fig. 5b). Across the population, drinking-epoch-selective neurons showed significant excitatory responses for the waiting and drinking epochs (red histogram at the top in Fig. 5b, p < 0.01, permutation test), and significant inhibitory responses for other behavioral epochs (blue histogram at the top in Fig. 5b, p < 0.01, permutation test). Alternatively, waiting-epoch-selective neurons showed significant excitatory responses for the late phase of go-cue odor-sampling and waiting epochs (red histogram at the bottom in Fig. 5b, p < 0.01, permutation test), and significant inhibitory responses for the drinking and no-go waiting epochs (blue histogram at the bottom in Fig. 5b, p < 0.01, permutation test). The waiting-epoch-selective neurons showed higher responses during the go-cue odor-sampling epoch than those of other groups (Fig. 5c, one-way analysis of variance with Tukey’s post hoc test). Thus, waiting-epoch-selective neurons exhibited associations between the go-cue excitations and excitatory responses for waiting behavior, suggesting that a subset of ACo neurons was involved in cue-outcome associations.

**Fig. 5.**
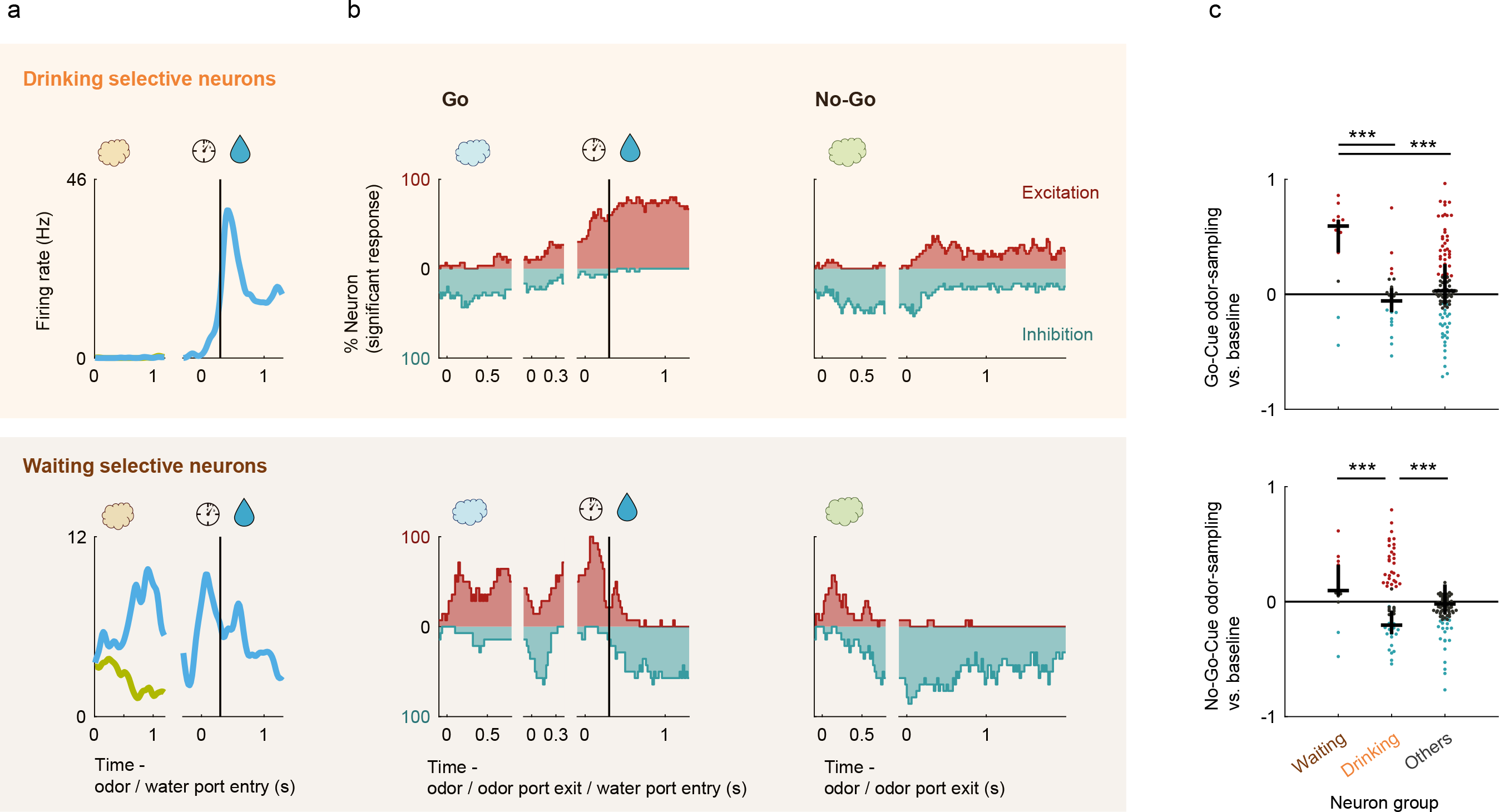
Waiting-selective neurons also showed go-cue odor-preferred responses during odor-sampling. (**a**) Example firing patterns of cue-outcome responses. Spike histograms are calculated using a 20 ms bin width and smoothed by convolving spike trains with a 60 ms wide Gaussian filter. A vertical black line indicates the water valve opening. (**b**) The proportions of neurons that exhibited significant excitatory and inhibitory response calculated from auROC values (p < 0.01, permutation test) for each neuron group. Vertical black lines indicate the water valve opening. (**c**) auROC values during odor-sampling epoch of go-cue odor-selective responses (top graph) and no-go-cue odor-selective responses (bottom graph) for each neuron group. Black horizontal lines and black vertical lines indicate medians and interquartile ranges. Red dots, significant excitation; blue dots, significant inhibition; gray dots, non-significant (p < 0.01, permutation test). Statistical significance among three groups (***P < 0.001) was assessed by one-way analysis of variance (ANOVA) with Tukey’s post hoc test.

## Discussion

The purpose of the study was to understand the electrophysiological features of ACo neurons on odor-evoked reward-related behavioral tasks. We found that many ACo neurons exhibited go-cue odor-preferred responses at the late phase of the odor-sampling epoch (Figs. 1-2). Consequently, the ACo population showed profound and persistent transformations in the dynamics of cue encoding, and provided sufficient information to account for the behavioral performance before executing the odor-evoked behaviors (Fig. 3). In addition to the late phase of odor-evoked activities, we also found two types of reward anticipation signals during the odor-evoked behaviors: ramp-like gradual increases in the signals to the reward exhibited by drinking-epoch-selective neurons, and phasic anticipation signals exhibited by waiting-epoch-selective-neurons (Fig. 4). The waiting-epoch-selective neurons exhibited associations between the go-cue excitations and excitatory responses for the waiting behavior (Fig. 5). Thus, the ACo showed unique encodings during various behavioral states in the task, suggesting that the ACo neurons play an important role in reward-related learning evoked by olfactory stimulus.

### Odor representation of the ACo

The ACo was a previously unexplored area in the olfactory amygdala located caudally to the lateral olfactory tract and rostromedially to the posterolateral cortical nucleus of the amygdala. The projection from the olfactory bulb (OB) terminates in the outer portion of the most superficial layer (layer Ia) of the cortex, and the projection from other olfactory cortex areas terminates in the deep portion of the ACo ^14,15^. Unlike other sensory systems, olfactory information from the external world reaches the amygdala without passing through thalamic areas. Therefore, the olfactory amygdala, including the ACo, receives olfactory information from the OB via the olfactory cortex ^16,17^. Moreover, previous comprehensive indirect evidence suggests that ACo may play a prominent role in reward-related behavioral learning from olfactory stimuli ^18,19^. However, little is known about the functions of the ACo in olfactory information processing. We performed *in vivo* recordings in the ACo neurons during the odor-guided go/no-go task to obtain a water reward. ACo neurons exhibited cue-odor-preferred responses at the late phase of odor-sampling epochs (Figs. 1,2, and 3). The peak firing in ACo neurons during odor-sampling epochs was later than that in other olfactory cortical areas (e.g., the piriform cortex ^20^; and the ventral tenia tecta ^21^). This late phase coding in the ACo neurons was not the premotor signal (Fig. 2). Population coding of ACo neurons showed profound and persistent transformations in the dynamics of cue-encoding, 400 ms after the odor onset (Fig. 3). This may reflect the input of other olfactory cortex from layers Ib, II, and III, and the top-down inputs from other brain areas, rather than the direct sensory inputs from OB in layer Ia. Therefore, we speculate that ACo neurons send task-modulated olfactory information to other amygdala areas.

### Role of reward-anticipation response in the waiting epoch

ACo sends the axons massively to the basolateral amygdala complex (BLA) ^3^. The BLA plays a role in learning and storing CS-US associations ^18,19^. Therefore, the projections from the ACo to the BLA may send olfactory information for the CS-US association in the odor-guided learning tasks. We demonstrated that some neurons exhibited reward-anticipation responses in the waiting epoch and also showed a go-cue-odor preference (Fig. 5). Since the ACo receives direct inputs from the BLA ^3^, it may also serve some function for the CS-US association by linking the olfactory information with the reward anticipation.

A previous study revealed that pyramidal neurons of the ACo displayed rhythmic fluctuations of the intrinsically generated voltage-dependent membrane potential in the theta-low beta range with the activation of sodium conductance ^9^. Synchronizing theta oscillations have been found to increase between regions when enhanced communication occurs during memory acquisition ^22,23^ and goal selection ^24,25^. Oscillatory synchronization for CS occurs in the theta band, between the lateral entorhinal cortex (LEC), which is a part of the olfactory cortex, and hippocampus (HPC), during the learning stage of trace CS-US associative learning tasks ^26^. The ACo has bidirectional connections with the LEC ^3^. We speculate that the ACo additionally drives the LEC-HPC circuit and supports the CS-US association by synchronizing the ACo-LEC-HPC theta oscillations during the learning stage.

### Role of reward-anticipation response in the drinking epoch

In learning, reward signals have very important implications ^27^. A subset of ACo neurons increased their firing rate during the drinking reward epoch (Fig. 4). These neurons started to increase their firing rate before the drinking epoch, and these activities persisted during the epoch. Previous studies reported that similar firing patterns were observed in the dopamine neurons in the VTA. Post learning, dopamine activity may change phasic responses to cues and rewards, and ramping activity may arise as the agent approaches the reward ^28^. The ACo receives direct inputs from the VTA ^3^. It is assumed that the ramping-like response in the ACo may reflect the inputs from the VTA. In addition, ACo has anatomical connections with other olfactory cortices ^3^. We speculate that the VTA reward signals may be transmitted to other olfactory cortical areas via ACo, making learning more efficient in the olfactory cortex.

A previous behavioral study revealed that electrical stimulation of the VTA showed that the ACo, besides other mesolimbic structures, displayed increased Fos expression in rats ^8^. ACo sends excitatory glutamatergic/aspartatergic projections to the nucleus accumbens (NAc) ^29^. Dopamine (DA) projections from the VTA to the NAc, which constitute the mesolimbic DA system ^30,31,32^, play an essential role in motivated behaviors, reinforcement learning, and reward processing ^33,34,35^. Therefore, the ACo may assist in driving the NAc-VTA circuit for reward-related behavior.

## Methods

### Animals

All experiments were performed on adult male C57BL/6 mice purchased from Shimizu Laboratory Supplies Co., Ltd., Kyoto, Japan (9 weeks old; weighing 20–25 g). The mice were individually housed in a temperature-controlled environment with a 13-hr light 11-hr dark cycle (lights on at 08:00 and off at 21:00). They were provided water after training and recording sessions so that body weights dipped no lower than 85% of initial levels and food was supplied ad libitum. All experiments were performed in accordance with the guidelines for animal experiments at Doshisha University and with the approval of the Doshisha University Animal Research Committee.

### Apparatus

We used a behavioral apparatus controlled by the Bpod State Machine r0.5 (Sanworks LLC, NY, USA), an open-source control device designed for behavioral tasks. The apparatus comprised of a custom-designed mouse behavior box with two nose-poke ports on the front wall. The box was contained in another soundproof box (BrainScience Idea. Co., Ltd., Osaka, Japan) equipped with a ventilator fan that provided adequate air circulation and low-level background noise. Each of the two nose-poke ports had a white light-emitting diode (LED) and an infrared photodiode. Interruption of the infrared beam generated a transistor-transistor-logic (TTL) pulse, thus signaling the entry of the mouse head into the port. The odor delivery port was equipped with a stainless steel tubing connected to a custom-made olfactometer ^36^. Eugenol was used as the go-cue odor and amyl acetate (Tokyo Chemical Industry Co., Ltd., Tokyo, Japan) as the no-go-cue odor respectively. These odors were diluted to 10% in mineral oil and further diluted 1:9 by airflow. Water-reward delivery was based on gravitational flow, controlled by a solenoid valve (The Lee Company, CT, USA), and connected via Tygon tubing to stainless steel tubing. The reward amount (6 μL) was determined by the opening duration of the solenoid valve, which was regularly calibrated.

### Odor-Guided go/no-go task

After a 3 s inter-trial interval, each trial began by illuminating the LED light at the right odor port, which instructed the mouse to poke its nose into that port. This resulted in the delivery of one of the two cue odors for 500 ms. Mice were required to keep their nose poked during odor stimulation to sniff the odor. After odor stimulation, the LED light was turned off and the mice could withdraw their nose from the odor port. If an eugenol odor (go-cue odor) was presented, the mice were required to move to the left water reward port and poke their nose within a timeout period of 2 s. At the water port, the mice were required to maintain their nose poke for 300 ms before water delivery began. Next, 6 μL of water was delivered as a reward. If an amyl acetate odor (no-go-cue odor) was presented, the mice were required to avoid entering the water port for 2 s following odor stimulation. Once in 10 trials, we introduced catch trials in which the air stream was delivered through a filter containing no odorants during which the mice were not rewarded regardless of their choice (go or no-go behavior). During the training sessions, mice learned to obtain water rewards at the left water port, move from the right odor port to the left odor port, and associate odor cues with the correct action. The accuracy rate was calculated as the total percentage of successes in the go and no-go trials in a session. The mice performed up to 524 trials (go error: ∼51 trials, no-go error: ∼13 trials, catch: ∼48 trials) in each session per day.

### Electrophysiology

Mice were anesthetized with medetomidine (0.75 mg/kg i.p.), midazolam (4.0 mg/kg i.p.), and butorphanol (5.0 mg/kg i.p.), and implanted with a custom-built microdrive of four tetrodes in the ACo (0.1 mm anterior to the bregma, 2.2 mm lateral to the midline). Individual tetrodes consisted of four twisted polyimide-coated tungsten wires (California Fine Wire, single wire diameter 12.5 μm, gold plated to less than 500 kΩ). Two additional screws were threaded into the bone above the cerebellum for reference. The electrodes were connected to an electrode interface board (EIB-18, Neuralynx, MT, USA) on a microdrive. The microdrive array was fixed to the skull using LOCTITE 454 (Henkel Corporation, Düsseldorf, Germany). After the completion of surgery, the mice received atipamezole (0.75 mg/kg i.p.) to reverse the effects of medetomidine and to allow for a shorter recovery period. The mice also received analgesics (ketoprofen, 5 mg/kg, i.p.). Behavioral training resumed at least 1 week after surgery. Electrical signals were obtained using open-source hardware (Open Ephys). For unit recordings, signals were sampled at 30 kHz in Open Ephys and band-pass filtered at 600–6,000 Hz. After each recording, the tetrodes were adjusted to obtain new units.

### Data analyses

All data analyses were performed using the built-in software in MATLAB 2019a (The Mathworks, Inc., MA, USA).

#### Spike sorting

Spikes were sorted into clusters offline using Kilosort2 (https://github.com/MouseLand/Kilosort2), with default parameters. Kilosort2 sorted spikes on the basis of spike waveform similarity, the bimodality of the distribution of waveform features, and the spike auto- and cross-correlograms. A unit was considered a single unit if Kilosort2 categorized that unit as “good.” Additional analyses and spike waveform plotting with data were performed with MATLAB code modified from N. Steinmetz (https://github.com/cortex-lab/spikes).To assess the quality of our recordings, we checked all spike waveforms defied by the “good” units with Kilosort 2, and some single units that had strange waveforms were excluded from the analyses.

#### Spike train analyses

Neural and behavioral data were synchronized by inputting each event timestamp from the Bpod behavioral control system into the electric signal recording system. For calculation of firing rates during tasks, peri-event time histograms (PETHs) were calculated using a 20 ms bin width, and smoothed by convolving spike trains with a 60 ms wide Gaussian filter.

#### ROC analyses

To quantify the firing rate changes, we used an algorithm, based on ROC analyses, that calculates the ability of an ideal observer to classify whether a given spike rate was recorded in one of two conditions (e.g., during go-cue or no-go-cue odor presentation) ^37^. We defined an auROC equal to 2 (ROCarea – 0.5), with the measure ranging from –1 to 1, where –1 signifies the strongest possible value for one alternative and 1 signifies the strongest possible value for the other.

The statistical significance of these ROC analyses was determined using a permutation test. For this test, we recalculated ROC curves after randomly reassigning all firing rates to either of the two groups arbitrarily, repeated this procedure a large number of times (500 repeats for analyses of dynamics [Figs. 1e, 4b-c and 5b], 1000 repeats for all other analyses [Fig. 5c]) to obtain a distribution of values. We then calculated the fraction of random values exceeding the actual value. For all analyses, we tested for significance at α = 0.01. Only neurons with a minimum number of three trials for each analyzed condition were included in the analyses.

For analyses of dynamics (width: 100 ms, step: 20 ms), we calculated three measures from the auROC values of correct trials (Figs. 4b and 4d):

1. Time of center of mass: the time corresponding to the center of mass of the significant points of the auROC values (p < 0.01, permutation test). Only neurons with significant points for each analyzed condition were included in this analysis.
2. Duration: The duration to the time of center of mass over which the auROC values were significant (p < 0.01, permutation test) for five or more consecutive bins, containing the time of center of mass. Only neurons with consecutive bins for each analyzed condition were included in this analysis.
3. Onset time: The time at which the duration was first evident.

#### Generalized linear models

To quantify the contribution of behavioral variables to neural activity, we used a generalized linear model (GLM), which was a multiple linear regression with the firing rate of each neuron as the dependent variable, and predictors derived from the behavioral variables as the independent variables (Fig. 2a and Supplementary Figs. 1a-c) ^13^. In this analysis, the firing rate (20 ms bin width and smoothed by convolving spike trains with a 60 ms wide Gaussian filter) of each neuron is described as a linear sum of temporal filters aligned to task events. For the current study, only odor stimulus onset, offset, and pre-odor port exit events were required, since we considered only the period between odor stimulus onset and 500 ms after the odor port exit. In the model, the predicted firing rate is given as:

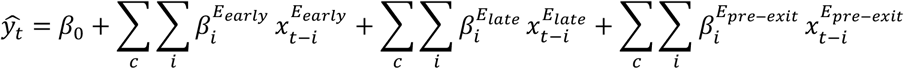

The response of a neuron at bin *t* is modeled 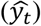 by the sum of a bias term (*β*_*0*_) and the weighted (*β*_*i*_) sum of various additional binary predictors at different lags (*i*), and *c* represents the two conditions (go or no-go trials). Binary predictors for the odor stimulus presentation 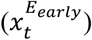 are supported over the window 0 to 500 ms relative to the onset of odor valve in either go or no-go trials (lags corresponding to odor presentation period, 25 time bins) as well as late phase of odor-sampling predictors 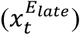 that are supported over the window 0 to 553/264 ms relative to the offset of the odor valve in either go or no-go trials (lags corresponding to the median durations between the odor valve offset and odor port exit, 28/14 time bins). Binary predictors for pre-odor port exit predictors 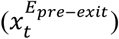 are supported over the window -300 to 0 ms relative to the odor port exit in either go or no-go trials (15 time bins). The *β* values were calculated using the glmfit MATLAB function.

Calculation of the relative contributions of behavioral variables to neural activity: We quantified the relative contribution of each behavioral variable to neural activity (Fig. 2b and Supplementary Figs. 1d-f) by determining how the performance of the encoding model declined when each variable was excluded from the model ^13,38^. We predicted the firing rate of each neuron in either case with all variables (full model), or by excluding one of the variables (partial model), with fivefold cross-validation (over trials; meaning that in each fold, 80% of trials were used for training the model and the remaining trials were used for testing the model performance). The relative contribution of each behavioral variable was calculated by comparing the variance explained by the partial model to the variance explained by the full model. For the current study, which included six behavioral variables, the relative contribution of each variable was defined as

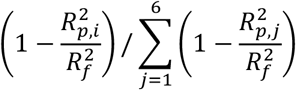

Here, 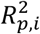 is the variance explained by the partial model that excludes the *i*th variable, and 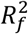is variance explained by the full model. Negative relative contributions were set to zero (this occurs when the R^2^ of the full model is lower than that of the partial model, owing to the introduction of noise by the excluded variable).

We used two approaches to exclude variables from the full model and calculated the variance explained by the partial model ^13^. In the first approach, the partial model was equivalent to the full model, except that the *β* values of the predictors of the excluded variable were set to zero (‘no refitting’). In the second approach, we calculated new *β* values by re-running the regression without the predictors of the excluded variable (refitting). Both approaches to exclude variables produced comparable results; the no-refitting approach was used to generate the main figures, and comparison with the refitting approach is shown in Supplementary Fig. 1d.

Moreover, we compared relative contributions as assessed separately using three different approaches: no refitting (NR; used in the paper), no refitting + Lasso regularization (NR + L), and refitting (R). Lasso regularization was applied using the lasso function in MATLAB; the mean square error (MSE) of the model was estimated using fivefold cross-validation, and we chose the lambda value that minimized the MSE. The results obtained with lasso regularization were almost identical to those obtained without regularization (Supplementary Fig. 1d), which suggested that there was no significant overfitting in our model.

Finally, to evaluate the significance of relative contributions assessed by the no-refitting approach, we calculated the control values. In this approach, the partial model was equivalent to the full model, except that the randomly selected *β* values of the predictors of the excluded variable (10% of predictors, mostly corresponding to the sum of time bins of each behavioral predictor) were set to zero, in which case, processing was performed 1,000 times (Supplementary Figs. 1e-f). Using the control mean ± 2 standard deviation (SD), the statistical significance was determined (< mean – 2SD, negative relative contribution; > mean + 2SD, positive relative contribution).

Population vector construction and analyses: We constructed 2 conditions (91 time bins) × 158 neurons matrix ^39,40,41^ during the odor-sampling epoch, in which columns contained the auROC values of the correct trials, corresponding to the trial-averaged firing rate changes from baseline (Supplementary Fig. 2a). By performing principal component analysis (PCA) on the dataset, we reduced the dimensionality of the ACo population from 158 neurons to three principal components (PCs), and obtained the odor-sampling epoch subspaces. Note that we used the three subspaces because they explained 80.6% of the total variance (Supplementary Fig. 2b). To visualize the ACo population responses, we projected the dataset onto three-dimensional subspaces (Fig. 3a). This allowed us to obtain a point reflecting the entire population response for each of the two conditions at a given instance. The distance between cue responses was computed as the Euclidean distance between pairs of activity vectors of all subspaces at a given instant (Fig. 3b) ^42,43^. This value was compared with the values during the baseline epoch (1200 to 1000 ms before the odor port entry).

#### SVM decoding analyses

We used a support vector machine (SVM) algorithm with a linear kernel as a classifier ^20,42^ and a MATLAB function (fitcsvm) for analyses. All analyses were conducted on trial data pooled across animals. A matrix containing concatenated firing rates for each trial was used, and each neuron was the input to the classifier. The matrix dimensions were the number of neurons by the number of trials. To avoid over-fitting, k-fold cross-validation (k = 10) was used to calculate the decoding accuracy of trial type discrimination. To compute decoding accuracy, forty trials for each trial type (from the start of the session) were chosen as the dataset. Next, the dataset was partitioned into ten equal parts; one part was used for testing, and the remaining parts were used for training the classifier. This process was repeated ten times to test each individual part, and the mean value of the accuracy was used for decoding accuracy. To compute the decoding accuracy of a 100 ms bin window (step: 20 ms), the classifier was trained and tested with a 100 ms bin window (step: 20 ms).

#### Statistical analyses

Data were analyzed using MATLAB 2019a. Statistical methods in each analysis are described above, in the result section, or in the figure legends. The Tukey-Kramer method was applied for significance tests with multiple comparisons. Sample sizes in this study were not pre-determined by calculation, they were based on previous research in the olfactory cortex fields ^20,44^. Randomization and blinding were not employed. Biological replicates for the histological studies are described in the figure legends.

### Histology

After recording, the mice were deeply anesthetized by intraperitoneal injection of sodium pentobarbital. Electric lesions were made using 10–20 μA direct current stimulation for 5 s of one of the four tetrode leads. Mice were perfused transcardially with phosphate-buffered saline (PBS) and 4% paraformaldehyde (PFA). Brains were removed from the skull and post-fixed in PFA. Brains were then cut into 50-μm-thick coronal sections and stained with cresyl violet. Electrode track positions were determined in reference to the atlas developed by Paxinos and Watson ^45^.

## Data availability

The data that support the findings of this study are available from the corresponding author upon reasonable request.

## Code availability

The custom code used for the analyses in the present study is available from the corresponding authors upon reasonable request.

## Acknowledgments

We thank Nozomi Fukui for assistance with data collection and the lab members for valuable discussions. We thank Hideki Tanisumi for providing illustrations in the figures. H.M. was supported by the Takeda Science Foundation and JSPS KAKENHI Grant Numbers 25135708 and 16K14557. Y.S. was supported by JSPS KAKENHI Grant Numbers 20H00109 and 20H05020.

## Author contributions

K.S., Y.T., and H.M. designed the experiments, and K.S., Y.T., and H.M. performed the experiments. K.S., Y.T., J.H., and H.M. performed the data analysis. K.S., Y.T., and H.M. wrote the paper. Y.S. supported and advised the project.

## Supplementary figure legends

**Supplementary Fig. 1.**
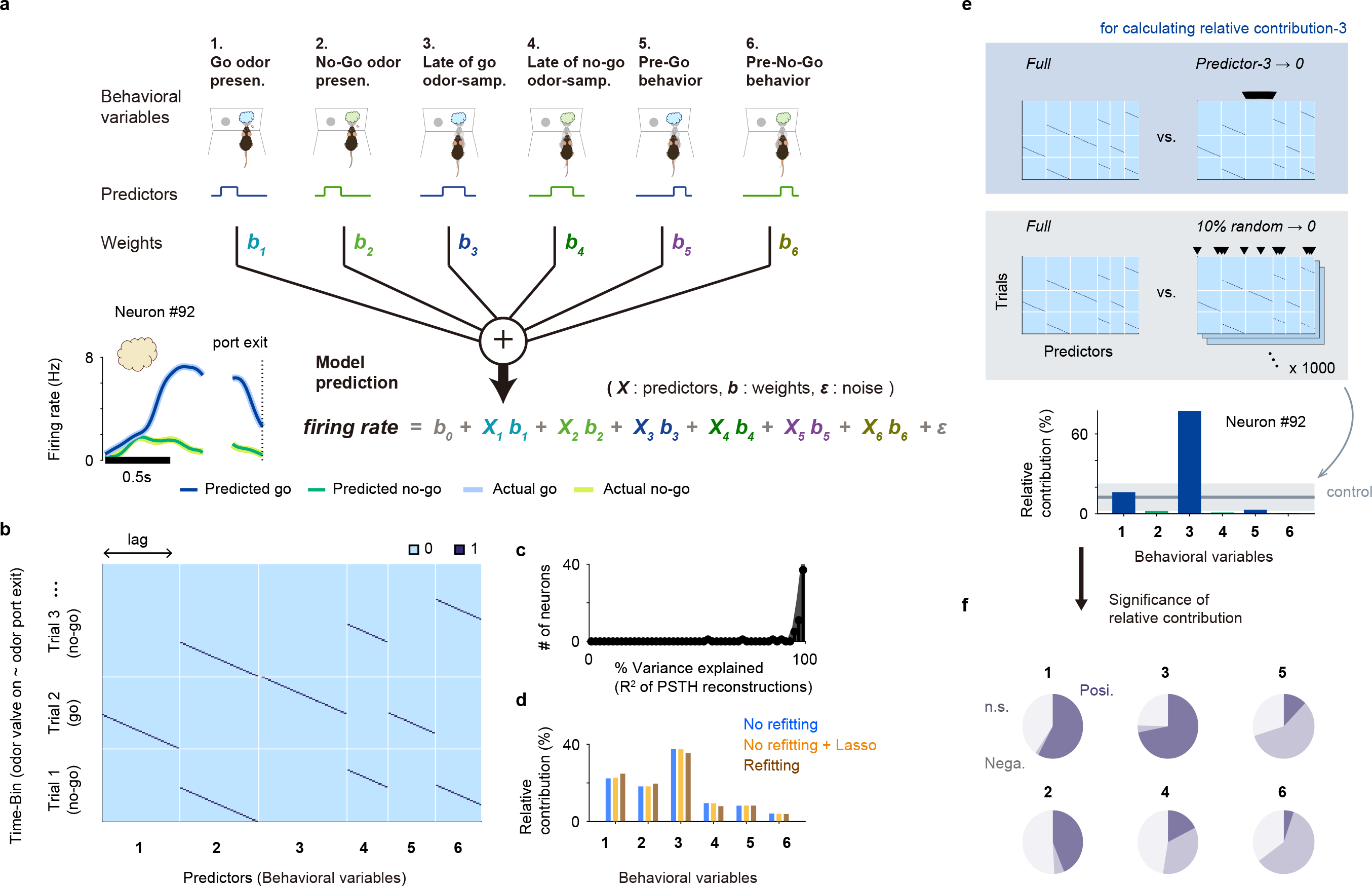
Generalized linear models and the relative contributions of behavioral variables to neural activity. (**a**) Schematic of the encoding model used to quantify the relationship between behavioral variables and the activity of each neuron. Inset, predicted and actual averaged firing rate relative to the odor onset and odor port exit for one neuron. (**b**) Structure of predictor matrices. The predictor has columns for each variable, which take non-zero values for time bins (rows) corresponding to the appropriate time offset from the given event. (**c**) Variance explained (R^2^ of PSTH reconstructions) between predicted and actual averaged firing rate relative to the odor onset and odor port exit across the go-cue-selective neurons. (**d**) Average relative contributions across the go-cue-selective neurons assessed separately using three different approaches: no refitting (used in the paper); no refitting + Lasso regularization; and refitting. Lasso regularization was applied using the lasso function in MATLAB; the mean square error (MSE) of the model was estimated using fivefold cross-validation, and we chose the lambda value that minimized the MSE. The results with lasso regularization were almost identical to the result without regularization, which suggests that there was no significant overfitting in our model. (**e**) Evaluation for significance of relative contributions assessed no refitting approach. The partial model was equivalent to the full model, except that the randomly selected *β* values of the predictors of the excluded variable (10% of predictors) were set to zero, in which processing was performed 1,000 times. Using the control mean ± 2 standard deviation (SD), the statistical significance was determined (< mean – 2SD, negative relative contribution; > mean + 2SD, positive relative contribution). (**f**) Proportions of the significance of relative contributions for each behavioral variable across the go-cue-selective neurons.

**Supplementary Fig. 2.**
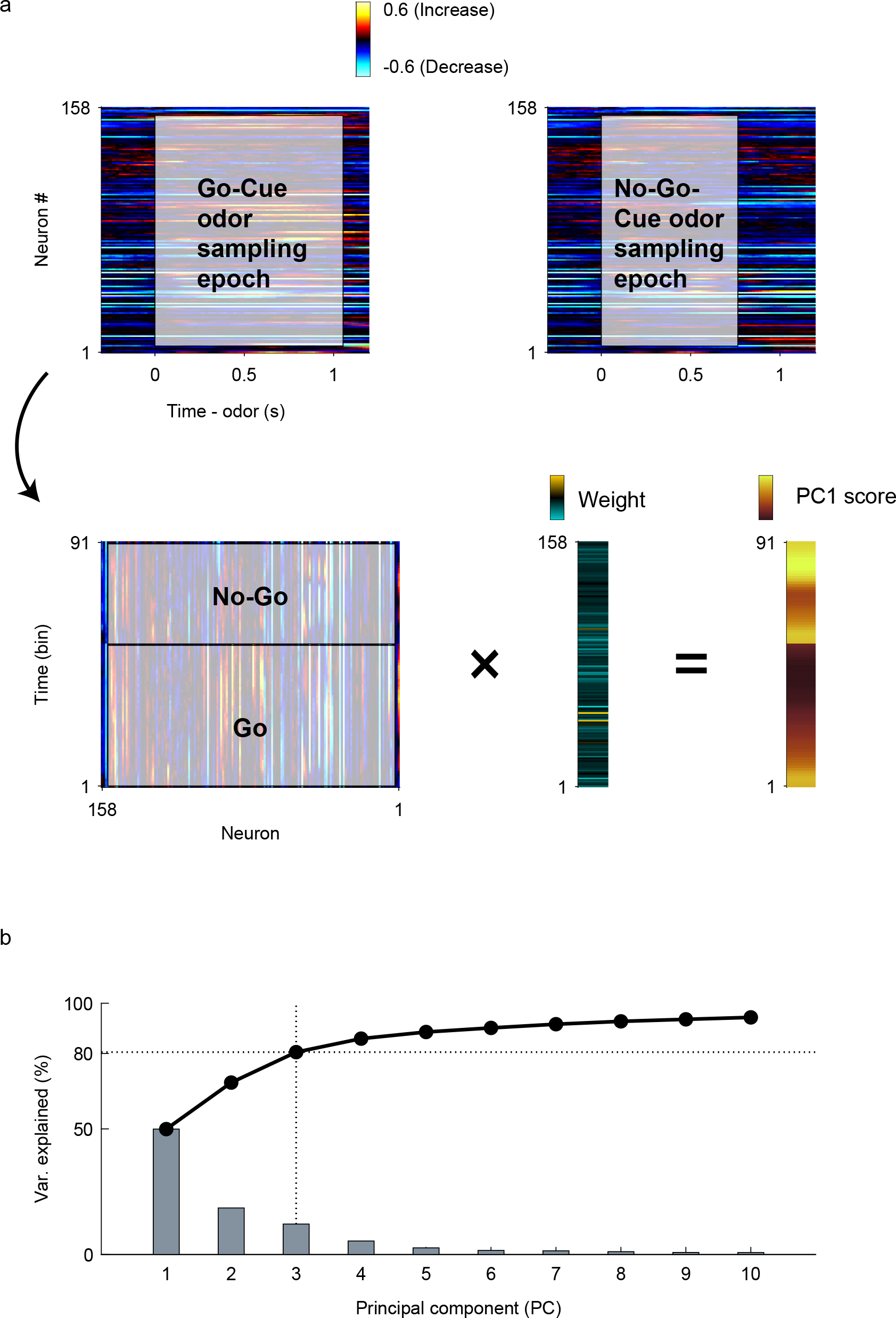
Population vector construction and analyses for ACo neuron population response. (**a**) Population vector construction. We constructed the two conditions (91 time bins) × 158 neurons matrix during the odor-sampling epoch, within which, the columns contained the auROC values corresponding to the trial-averaged firing rate changes from the baseline. By performing principal component analysis (PCA) on the dataset, we reduced the dimensionality of the ACo population from 158 neurons to three principal components (PCs). Subsequently, we obtained the odor-sampling epoch subspaces (graphs show the values of the first dimension of the odor-sampling epoch subspaces). (**b**) Screen plot of the odor-sampling epoch subspaces. It is notable that we used the three subspaces because they explained 80.6% of the total variance.

## References

1. Ehrlichman, H. & Bastone, L. Olfaction and Emotion BT - Science of Olfaction. in (eds Serby, M. J. & Chobor, K. L.) 410–438 (Springer New York, 1992). doi:10.1007/978-1-4612-2836-3_15

2. Soudry, Y., Lemogne, C., Malinvaud, D., Consoli, S.-M. & Bonfils, P. Olfactory system and emotion: Common substrates. Eur. Ann. Otorhinolaryngol. Head Neck Dis. 128, 18–23 (2011).

3. Cádiz-Moretti, B., Abellán-Álvaro, M., Pardo-Bellver, C., Martínez-García, F. & Lanuza, E. Afferent and efferent projections of the anterior cortical amygdaloid nucleus in the mouse. J. Comp. Neurol. 525, 2929–2954 (2017).

4. Zhou, G., Lane, G., Cooper, S. L., Kahnt, T. & Zelano, C. Characterizing functional pathways of the human olfactory system. Elife 8, (2019).

5. Haberly, L. B. Parallel-distributed Processing in Olfactory Cortex: New Insights from Morphological and Physiological Analysis of Neuronal Circuitry. Chem. Senses 26, 551–576 (2001).

6. Mori, K. & Sakano, H. How is the olfactory map formed and interpreted in the mammalian brain? Annu Rev Neurosci 34, 467–499 (2011).

7. Sevelinges, Y., Gervais, R., Messaoudi, B., Granjon, L. & Mouly, A.-M. Olfactory fear conditioning induces field potential potentiation in rat olfactory cortex and amygdala. Learn. Mem. 11, 761–769 (2004).

8. Majkutewicz, I. et al.. Lesion of the ventral tegmental area amplifies stimulation-induced Fos expression in the rat brain. Brain Res. 1320, 95–105 (2010).

9. Sanhueza, M. & Bacigalupo, J. Intrinsic subthreshold oscillations of the membrane potential in pyramidal neurons of the olfactory amygdala. Eur. J. Neurosci. 22, 1618–1626 (2005).

10. Steinmetz, N. A., Zatka-Haas, P., Carandini, M. & Harris, K. D. Distributed coding of choice, action and engagement across the mouse brain. Nature 576, 266–273 (2019).

11. Allen, W. E. et al.. Thirst regulates motivated behavior through modulation of brainwide neural population dynamics. Science (80-.). 364, eaav3932 (2019).

12. Parker, P. R. L., Brown, M. A., Smear, M. C. & Niell, C. M. Movement-Related Signals in Sensory Areas: Roles in Natural Behavior. Trends Neurosci. 43, 581–595 (2020).

13. Engelhard, B. et al. Specialized coding of sensory, motor and cognitive variables in VTA dopamine neurons. Nature 570, 509–513 (2019).

14. Scalia, F. & Winans, S. S. The differential projections of the olfactory bulb and accessory olfactory bulb in mammals. J. Comp. Neurol. 161, 31–55 (1975).

15. Carmichael, S. T., Clugnet, M. C. & Price, J. L. Central olfactory connections in the macaque monkey. J. Comp. Neurol. 346, 403–434 (1994).

16. Price, J. L. An autoradiographic study of complementary laminar patterns of termination of afferent fibers to the olfactory cortex. J. Comp. Neurol. 150, 87–108 (1973).

17. McDonald, A. J. Cortical pathways to the mammalian amygdala. Prog. Neurobiol. 55, 257–332 (1998).

18. Adhikari, A. et al. Basomedial amygdala mediates top-down control of anxiety and fear. Nature 527, 179–185 (2015).

19. Gore, F. et al. Neural Representations of Unconditioned Stimuli in Basolateral Amygdala Mediate Innate and Learned Responses. Cell 162, 134–145 (2015).

20. Miura, K., Mainen, Z. F. & Uchida, N. Odor representations in olfactory cortex: distributed rate coding and decorrelated population activity. Neuron 74, 1087–1098 (2012).

21. Shiotani, K. et al. Tuning of olfactory cortex ventral tenia tecta neurons to distinct task elements of goal-directed behavior. Elife 9, (2020).

22. Hoffmann, L. C. & Berry, S. D. Cerebellar theta oscillations are synchronized during hippocampal theta-contingent trace conditioning. Proc. Natl. Acad. Sci. U. S. A. 106, 21371–21376 (2009).

23. Wikgren, J., Nokia, M. S. & Penttonen, M. Hippocampo-cerebellar theta band phase synchrony in rabbits. Neuroscience 165, 1538–1545 (2010).

24. Jones, M. W. & Wilson, M. A. Theta rhythms coordinate hippocampal-prefrontal interactions in a spatial memory task. PLoS Biol. 3, e402 (2005).

25. Benchenane, K. et al. Coherent theta oscillations and reorganization of spike timing in the hippocampal-prefrontal network upon learning. Neuron 66, 921–936 (2010).

26. Takehara-Nishiuchi, K., Maal-Bared, G. & Morrissey, M. D. Increased Entorhinal-Prefrontal Theta Synchronization Parallels Decreased Entorhinal-Hippocampal Theta Synchronization during Learning and Consolidation of Associative Memory. Front. Behav. Neurosci. 5, 90 (2011).

27. Glimcher, P. W. Understanding dopamine and reinforcement learning: the dopamine reward prediction error hypothesis. Proc. Natl. Acad. Sci. U. S. A. 108 Suppl, 15647–15654 (2011).

28. Song, M. R. & Lee, S. W. Dynamic resource allocation during reinforcement learning accounts for ramping and phasic dopamine activity. Neural Netw. 126, 95–107 (2020).

29. Ray, J. P., Russchen, F. T., Fuller, T. A. & Price, J. L. Sources of presumptive glutamatergic/aspartatergic afferents to the mediodorsal nucleus of the thalamus in the rat. J. Comp. Neurol. 320, 435–456 (1992).

30. Björklund, A. & Dunnett, S. B. Dopamine neuron systems in the brain: an update. Trends Neurosci. 30, 194–202 (2007).

31. Ikemoto, S. Dopamine reward circuitry: two projection systems from the ventral midbrain to the nucleus accumbens-olfactory tubercle complex. Brain Res. Rev. 56, 27–78 (2007).

32. Morales, M. & Margolis, E. B. Ventral tegmental area: cellular heterogeneity, connectivity and behaviour. Nat. Rev. Neurosci. 18, 73–85 (2017).

33. Hamid, A. A. et al. Mesolimbic dopamine signals the value of work. Nat. Neurosci. 19, 117–126 (2016).

34. Salamone, J. D. & Correa, M. The mysterious motivational functions of mesolimbic dopamine. Neuron 76, 470–485 (2012).

35. Watabe-Uchida, M., Eshel, N. & Uchida, N. Neural Circuitry of Reward Prediction Error. Annu. Rev. Neurosci. 40, 373–394 (2017).

36. Uchida, N. & Mainen, Z. F. Speed and accuracy of olfactory discrimination in the rat. Nat. Neurosci. 6, 1224–1229 (2003).

37. Felsen, G. & Mainen, Z. F. Neural Substrates of Sensory-Guided Locomotor Decisions in the Rat Superior Colliculus. Neuron 60, 137–148 (2008).

38. Pho, G. N., Goard, M. J., Woodson, J., Crawford, B. & Sur, M. Task-dependent representations of stimulus and choice in mouse parietal cortex. Nat. Commun. 9, 2596 (2018).

39. Murray, J. D. et al. Stable population coding for working memory coexists with heterogeneous neural dynamics in prefrontal cortex. Proc. Natl. Acad. Sci. U. S. A. 114, 394–399 (2017).

40. Cavanagh, S. E., Towers, J. P., Wallis, J. D., Hunt, L. T. & Kennerley, S. W. Reconciling persistent and dynamic hypotheses of working memory coding in prefrontal cortex. Nat. Commun. 9, 3498 (2018).

41. Ohnuki, T., Osako, Y., Manabe, H., Sakurai, Y. & Hirokawa, J. Dynamic coordination of the perirhinal cortical neurons supports coherent representations between task epochs. Commun. Biol. 3, 406 (2020).

42. Cury, K. M. & Uchida, N. Robust odor coding via inhalation-coupled transient activity in the mammalian olfactory bulb. Neuron 68, 570–585 (2010).

43. Mazor, O. & Laurent, G. Transient dynamics versus fixed points in odor representations by locust antennal lobe projection neurons. Neuron 48, 661–673 (2005).

44. Manabe, H., Kusumoto-Yoshida, I., Ota, M. & Mori, K. Olfactory cortex generates synchronized top-down inputs to the olfactory bulb during slow-wave sleep. J. Neurosci. 31, 8123–8133 (2011).

45. Paxinos, G. The mouse brain in stereotaxic coordinates / George Paxinos, Keith Franklin. (Academic, 2004).

